# HIV coinfection is associated with low fitness *rpoB* variants in rifampicin-resistant *Mycobacterium tuberculosis*

**DOI:** 10.1101/2020.07.20.208835

**Authors:** Chloé Loiseau, Daniela Brites, Miriam Reinhard, Kathrin Zürcher, Sonia Borrell, Marie Ballif, Lukas Fenner, Helen Cox, Liliana K. Rutaihwa, Robert J. Wilkinson, Marcel Yotebieng, E Jane Carter, Alash’le Abimiku, Olivier Marcy, Eduardo Gotuzzo, Anchalee Avihingsanon, Nicola Zetola, Basra Doulla, Erik C. Böttger, Matthias Egger, Sebastien Gagneux

**Affiliations:** Swiss Tropical and Public Health Institute, Basel, Switzerland; University of Basel, Basel, Switzerland; Institute of Social and Preventive Medicine, University of Bern, Bern, Switzerland; Institute of Infectious Disease and Molecular Medicine and Wellcome Centre for Infectious Disease Research in Africa, University of Cape Town, Cape Town, South Africa; Ifakara Health Institute, Bagamoyo, Tanzania; Wellcome Centre for Infectious Diseases Research in Africa, University of Cape Town, Observatory 7925, South Africa; Department of Infectious Diseases, Imperial College London, London, UK; Francis Crick Institute, London NW1 1AT, UK; Division of General Internal Medicine, Department of Medicine, Albert Einstein College of Medicine, Bronx, NY, USA; Department of Medicine, Moi University School of Medicine, and Moi Teaching and Referral Hospital, Eldoret, Kenya; Institute of Human Virology, Abuja, Nigeria; Centre de Prise en Charge de Recherche et de Formation, Yopougon, Abidjan, Côte d’Ivoire; Bordeaux Population Health Research Center, Inserm U1219, University of Bordeaux, Bordeaux, France; TB Research Unit, Instituto de Medicina Tropical Alexander von Humboldt, Universidad Peruana Cayetano Heredia, Lima, Peru; The HIV Netherlands Australia Thailand (HIV-NAT) Research Collaboration, Thai Red Cross AIDS Research Centre and Tuberculosis research unit, Faculty of Medicine, Chulalongkorn University, Bangkok, Thailand; University of Pennsylvania, Philadelphia, PA, USA; Central Tuberculosis Reference Laboratory, Dar es Salaam, Tanzania; National Tuberculosis and Leprosy Programme, Dar es Salaam, Tanzania; Institute of Medical Microbiology, University of Zurich, Zurich, Switzerland; Swiss National Center for Mycobacteria, Zurich, Switzerland; Centre for Infectious Disease Epidemiology and Research, Faculty of Health Sciences, University of Cape Town, Cape Town, South Africa

## Abstract

We analysed 312 drug-resistant genomes of *Mycobacterium tuberculosis (Mtb)* collected from HIV coinfected and HIV negative TB patients from nine countries with a high tuberculosis burden. We found that rifampicin-resistant *Mtb* strains isolated from HIV coinfected patients carried disproportionally more resistance-conferring mutations in *rpoB* that are associated with a low fitness in the absence of the drug, suggesting these low fitness *rpoB* variants can thrive in the context of reduced host immunity.

---

Tuberculosis (TB), caused by members of the *Mycobacterium tuberculosis* (*Mtb*) Complex, is a leading cause of death worldwide, killing more people than any other infectious disease. Among the many factors driving the global TB epidemics, two factors stand out as particularly important: antibiotic resistance and HIV coinfection (1). Although the impact of both of these factors individually is well recognized, the interaction between them is less clear and likely depends on the particular epidemiologic setting (2). HIV coinfection and drug-resistant TB often coexist in severe epidemics, which could indicate spread of drug-resistant *Mtb* strains from immune-compromised patients (3–5). The propensity of drug-resistant *Mtb* strains to spread is influenced by the fitness cost associated with drug resistance determinants (6). Specifically, bacterial strains that have acquired drug resistance-conferring mutations may be less transmissible than their susceptible counterparts, although this fitness cost can be ameliorated by compensatory mutations (7–10). Moreover, the effect of different resistance-conferring mutations on fitness can be heterogeneous (11). In the clinical setting, there is a selection for high-fitness and/or compensated drug-resistant *Mtb* strains in TB patients (12). However, in immune-compromised hosts, such as HIV coinfected patients, even strains with low-fitness resistance mutations might propagate efficiently (13–15), which could partially explain why drug-resistant TB has been associated with HIV co-infection (16, 17). However, to date, no evidence directly supports the notion that the immunological environment created by HIV co-infection modifies the fitness of drug-resistant *Mtb* (5, 18, 19).

In this study, we tested the hypothesis that resistance-conferring mutations with low fitness in *Mtb* are overrepresented among HIV-coinfected TB patients. We focused our analysis on isoniazid and rifampicin, the two most important first-line anti-TB drugs, for which resistance-conferring mutations have been shown to differ in their fitness effects when measured in the laboratory (11). In addition, the frequency of the resistance alleles found in a clinical setting correlates well with the *in vitro* fitness of strains (20, 21). To explore the association between HIV coinfection and the fitness effect of different drug resistance-conferring mutations in *Mtb*, we compiled a collection of drug-resistant strains using the global International epidemiology Databases to Evaluate AIDS (IeDEA, http://www.iedea.org) consortium (22, 23) as a platform. For this study, 312 strains were collected from HIV-coinfected and HIV uninfected TB patients originating from nine countries on three continents: Peru, Thailand, South Africa, Kenya, Côte d’Ivoire, Botswana, Democratic Republic of the Congo, Nigeria and Tanzania (supplemental methods, Figure 1 and supplemental Table S1). The association between the fitness of isoniazid resistance-conferring mutations and HIV coinfection was tested in a univariate analysis (Figure S1). Isoniazid resistance-conferring mutations were divided into three groups, as previously described (24): *katG* S315T mutation, *katG* mutations other than S315T, and *inhA* promoter mutations only. The S315T substitution in *katG* causes high-level isoniazid resistance, while retaining some catalase/peroxidase functions (25). Conversely, the *inhA* promoter mutation does not affect KatG activity. Other substitutions/deletions in *katG* have been associated with a lower fitness in the laboratory and are observed only rarely among clinical isolates (24, 26, 27). In the case of rifampicin, the association between the fitness of *rpoB* variants and HIV coinfection was tested in both a univariate and multivariate analysis (Table 1). Resistance-conferring variants in *rpoB* were classified into two groups based on their fitness effects documented previously (11, 28, 29). The mutation *rpoB* S450L was considered ‘high-fitness’, since this mutation was previously shown to confer a low fitness cost in the laboratory (11) and is generally the most common in clinical strains (30). Any other resistance-conferring mutation affecting *rpoB* was considered ‘low-fitness’ (11). The multivariable logistic regression model with outcome “low fitness *rpoB* variants” was adjusted for host-related factors (history of TB, country of isolation, sex and age) (31) and bacterial factors (*Mtb* lineage, presence of a *rpoA/C* compensatory mutation, clustering of the genome inferred by genetic relatedness). Seventy-six patients from Tanzania and Botswana were excluded from the model due to missing or unknown clinical data.

**Table 1.** Results of the univariate and multivariate analysis showing host and bacterial factors associated with low fitness *rpoB* variants in 203 TB patients.

| Dependent:<br>fitness of *rpoB* variants |  | Low-fitness | High-fitness | univariable |  | multivariable |  |
| --- | --- | --- | --- | --- | --- | --- | --- |
|  |  |  |  | OR (95% CI) | *P* | OR (95% CI) | *P* |
| HIV status | HIV- | 71 (51.4) | 67 (48.6) | reference |  | reference |  |
|  | HIV+ | 47 (72.3) | 18 (27.7) | 2.46 (1.30-4.66) | 0.006 | 4.58 (1.69-12.44) | 0.003 |
| Presence of a compensatory mutation in *rpoA/C* | No | 117 (71.3) | 47 (28.7) | reference |  | reference |  |
|  | Yes | 1 (2.6) | 38 (97.4) | 0.01 (0.00-0.08) | < 0.0001 | 0.01 (0.00-0.06) | < 0.0001 |
| *Mtb* lineage | Lineage 2 | 16 (44.4) | 20 (55.6) | reference |  | reference |  |
|  | Lineage 4 | 99 (61.5) | 62 (38.5) | 2.00 (0.96-4.14) | 0.06 | 3.10 (0.94-10.21) | 0.06 |
|  | Other (L1 or L3) | 3 (50.0) | 3 (50.0) | 1.25 (0.22-7.05) | 0.80 | 0.97 (0.11-8.31) | 0.98 |
| Clustering of the genome | No | 109 (59.6) | 74 (40.4) | reference |  | reference |  |
|  | Yes | 9 (45.0) | 11 (55.0) | 0.56 (0.22-1.41) | 0.21 | 1.05 (0.28-3.90) | 0.94 |
| Country of isolation | South Africa | 29 (55.8) | 23 (44.2) | reference |  | reference |  |
|  | Democratic Republic of Congo | 11 (37.9) | 18 (62.1) | 0.48 (0.19-1.23) | 0.13 | 0.39 (0.12-1.34) | 0.14 |
|  | Côte d'Ivoire | 35 (79.5) | 9 (20.5) | 3.08 (1.24-7.70) | 0.02 | 2.04 (0.58-7.23) | 0.27 |
|  | Kenya | 4 (66.7) | 2 (33.3) | 1.59 (0.27-9.44) | 0.61 | 0.94 (0.10-8.42) | 0.96 |
|  | Nigeria | 20 (58.8) | 14 (41.2) | 1.13 (0.47-2.72) | 0.78 | 1.00 (0.29-3.40) | 0.99 |
|  | Peru | 16 (53.3) | 14 (46.7) | 0.91 (0.37-2.23) | 0.83 | 1.49 (0.33-6.70) | 0.60 |
|  | Thailand | 3 (37.5) | 5 (62.5) | 0.48 (0.10-2.20) | 0.34 | 0.42 (0.07-2.65) | 0.36 |
| Age | Mean (SD) | 32.5 (10.4) | 34.3 (12.3) | 0.99 (0.96-1.01) | 0.25 | 0.97 (0.94-1.01) | 0.10 |
| Sex | Female | 47 (59.5) | 32 (40.5) | reference |  |  |  |
|  | Male | 71 (57.3) | 53 (42.7) | 0.91 (0.51-1.62) | 0.75 | 0.77 (0.34-1.71) | 0.52 |
| History of TB disease | No | 35 (52.2) | 32 (47.8) | reference |  |  |  |
|  | Yes | 83 (61.0) | 53 (39.0) | 1.43 (0.79-2.58) | 0.23 | 0.96 (0.34-2.73) | 0.94 |

Number of observations in model = 203; CI = confidence interval; The odds ratio and p-value are obtained from the regression model.

**Figure 1.**
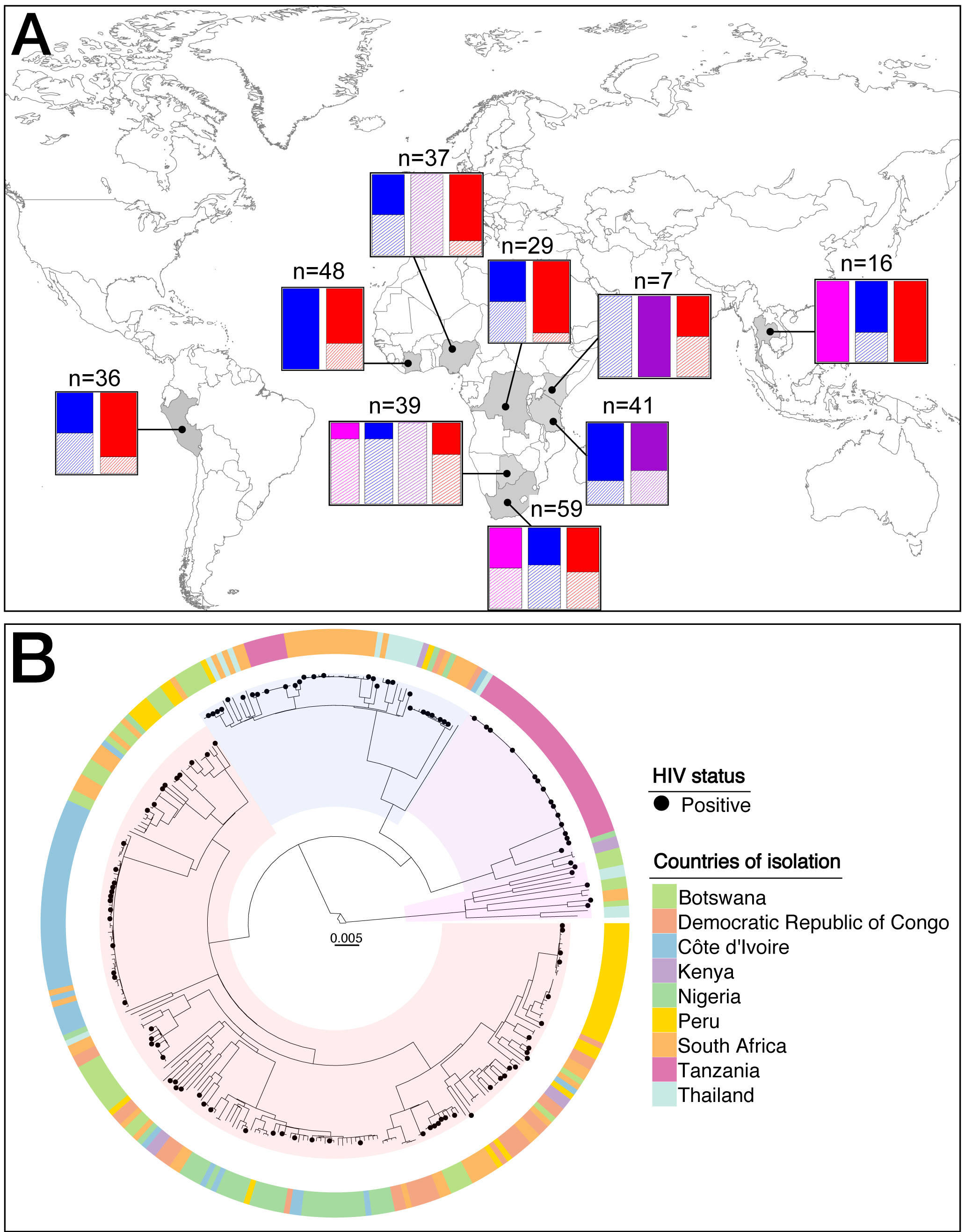
**A. Frequency of *Mtb* lineages by HIV status for countries sampled.** Countries coloured in grey were sampled. The barplots indicate the proportion of each lineage represented in this study. Magenta corresponds to *Mtb* lineage 1, blue corresponds to *Mtb* lineage 2, purple corresponds to *Mtb* lineage 3 and red corresponds to *Mtb* lineage 4. Solid colour corresponds to HIV negative and hatches correspond to HIV coinfected TB patients. The number of genomes sampled in each country is indicated on top of the barplots. **B. Phylogenetic tree of the dataset used in the study.** Maximum likelihood phylogeny of 312 whole-genome sequences based on 18,531 variable positions. The scale bar indicates the number of substitutions per polymorphic site. The phylogeny was rooted on *M. canettii. Mtb* isolated from HIV coinfected patients are indicated by black dots. The peripheral ring depicts the country of isolation of the strains sequenced.

Out of 312 patients, 113 (36.2%) were HIV-coinfected, 120 (38.5%) were women, 115 (37%) were newly diagnosed TB cases (therefore treatment naïve), 276 (88.5%) harboured isoniazid resistance-conferring mutations, with or without additional resistance, and 282 (90.4%) harboured rifampicin resistance-conferring mutations, with or without additional resistance. In total, 78.8% (n= 246) of the strains were classified as being at least MDR, defined as resistance to isoniazid and rifampicin with or without additional resistance to 2^nd^ line drugs. Amongst the 113 HIV coinfected individuals, 34 (30%) were on antiretroviral therapy (ART), 26 (23%) were not, and 53 (47%) had unknown ART start date. Four of the seven known *Mtb* lineages were represented in the following proportions: 11 L1 (3.5%), 57 L2 (18.3%), 38 L3 (12.2%), 206 L4 (66.0%). After dividing a total of 276 isoniazid-resistant strains into the three groups of isoniazid resistance-conferring mutations defined above, we found similar proportions in HIV-coinfected and HIV-uninfected patients (chi-square test, p=0.54, Figure S1), and as expected, the *katG* S315T mutation was the most frequent mutation in both categories (overall, found in 80% of isoniazid-resistant strains). In the case of rifampicin resistance, a univariate and multivariate analysis of 203 strains with complete clinical records, indicated that HIV coinfected TB patients carried a higher proportion of low-fitness *rpoB* resistance variants in comparison to HIV negative patients (72.3% vs. 51.4%). The univariate analysis showed higher odds of having a low-fitness *rpoB* variant in HIV coinfected patients (Odds Ratio 2.46 [95% Confidence Interval 1.30-4.66], p=0.006, Table 1). Our multivariable regression analysis confirmed these results and showed an association between low-fitness *rpoB* variants and HIV-coinfection, whilst controlling for other factors (Odds Ratio 4.58 [95% Confidence Interval 1.69, 12.44], p=0.003, Table 1). This association can be explained at least in two ways. Firstly, HIV-coinfected patients are thought to have fewer lung cavities on average and lower sputum bacillary load (32, 33). The resulting smaller *Mtb* population size would lead to fewer replication events, possibly reducing the number of mutations available for selection to act upon. In other words, low-fitness variants and high-fitness variants would co-occur less often in an HIV-coinfected patient, such that competition between them would be less likely. This scenario would be relevant for *de novo* acquisition of low-fitness drug-resistant variants within an HIV-coinfected patient. Secondly, following the transmission of a drug-resistant strain with low fitness to a host with reduced immunity, weaker immune pressure acting on this strain might lead to better bacterial survival. The association between low-fitness *rpoB* variants and HIV coinfection remained significant even after adjusting for the different epidemiologic settings (i.e. countries) and the strain genetic background (i.e. *Mtb* lineages). We also observed that strains carrying the *rpoB* S450L resistance-conferring mutation were more likely to also carry a compensatory mutation in *rpoA/C* (97.4% vs. 2.6%, Table 1). Even though this phenomenon seems counter-intuitive, it has been described multiple times (7, 9, 34–36) and might thus point to different mechanisms of compensation in strains carrying resistance mutations other than *rpoB* S450L. In addition, in our study, L4 strains were associated with low fitness *rpoB* variants, compared to L2 (Odds Ratio 3.10 [95% Confidence Interval 0.94, 10.21], p=0.06, Table 1), indicating that the strain genetic background could play a role in shaping the cost of resistance, as was previously shown for other bacterial species (37) and for other drugs (38). In the regression analysis, we had several categorical variables with only few observations. Therefore, statistical power especially for country of isolation was low and the results should be interpreted with care.

HIV coinfected TB patients are generally thought of having a reduced potential for TB transmission (32, 39) because these patients have reduced formation of lung cavities, more extrapulmonary disease, and a shorter period of infectiousness due to earlier diagnosis or higher mortality, especially in the absence of anti-retroviral treatment and if antibiotic resistance is already present (4). Based on the over-representation of low-fitness *rpoB* mutations in the context of HIV coinfection, one would expect a further reduction of the transmission potential of drug-resistant TB in this context. Yet, outbreaks of drug-resistant TB in HIV coinfected patients have been reported (40). Such outbreaks might be explained by i) a higher risk of *Mtb* infection and reinfection due to diminished host immunity, ii) on-going transmission of drug-resistant *Mtb* from a larger pool of immune-competent TB patients to immune-compromised patients, iii) transmission occurring in conducive environments such as health care settings where both HIV coinfected individuals and DR-TB patients are more likely to co-exist, and iv) *Mtb* strains carrying high-fitness drug resistance mutations.

In summary, using a global sample of drug-resistant *Mtb* clinical strains from HIV coinfected and HIV negative TB patients, we showed that low-fitness *rpoB* variants were overrepresented in HIV coinfected patients, and that this association was independent from other potential confounding factors. Taken together, our results provide new insights into how HIV coinfection can impact the fitness of drug-resistant *Mtb*.

## Data availability

The *Mtb* whole-genome sequences from the patients are available on NCBI under several project IDs. The accession number for each genome is indicated in the supplemental Table 1.

## Acknowledgments

We thank all the sites that participated in this study and the patients whose data were used in this study. We thank Dr. Jan Hattendorf for providing statistical help and Dr. Sebastian M. Gygli for critically reading the manuscript. Calculations were performed at the sciCORE (http://scicore.unibas.ch/) scientific computing core facility at University of Basel.

## References

1. World Health Organization. 2019. Global Tuberculosis Report.

2. Getahun H, Gunneberg C, Granich R, Nunn P. 2010. HIV Infection–Associated Tuberculosis: The Epidemiology and the Response. Clin Infect Dis 50:S201–S207.

3. Wells CD, Cegielski JP, Nelson LJ, Laserson KF, Holtz TH, Finlay A, Castro KG, Weyer K. 2007. HIV Infection and Multidrug-Resistant Tuberculosis--The Perfect Storm. J Infect Dis 196:S86–107.

4. Gandhi NR, Moll A, Sturm AW, Pawinski R, Govender T, Lalloo U, Zeller K, Andrews J, Friedland G. 2006. Extensively drug-resistant tuberculosis as a cause of death in patients co-infected with tuberculosis and HIV in a rural area of South Africa. Lancet 368:1575–1580.

5. Eldholm V, Rieux A, Monteserin J, Lopez JM, Palmero D, Lopez B, Ritacco V, Didelot X, Balloux F. 2016. Impact of HIV co-infection on the evolution and transmission of multidrug-resistant tuberculosis. Elife 5:1–19.

6. Cohen T, Murray M. 2004. Modeling epidemics of multidrug-resistant M. tuberculosis of heterogeneous fitness. Nat Med 10(10):1117–21.

7. Comas I, Borrell S, Roetzer A, Rose G, Malla B, Kato-Maeda M, Galagan J, Niemann S, Gagneux S. 2012. Whole-genome sequencing of rifampicin-resistant Mycobacterium tuberculosis strains identifies compensatory mutations in RNA polymerase genes. Nat Genet 44:106–10.

8. Casali N, Nikolayevskyy V, Balabanova Y, Ignatyeva O, Kontsevaya I, Harris SR, Bentley SD, Parkhill J, Nejentsev S, Hoffner SE, Horstmann RD, Brown T, Drobniewski F. 2012. Microevolution of extensively drug-resistant tuberculosis in Russia. Genome Res 22:735–45.

9. De Vos M, Müller B, Borrell S, Black PA, Van Helden PD, Warren RM, Gagneux S, Victor TC. 2013. Putative compensatory mutations in the rpoc gene of rifampin-resistant mycobacterium tuberculosis are associated with ongoing transmission. Antimicrob Agents Chemother 57:827–832.

10. Shcherbakov D, Akbergenov R, Matt T, Sander P, Andersson DI, Böttger EC. 2010. Directed mutagenesis of mycobacterium smegmatis 16S rRNA to reconstruct the in vivo evolution of aminoglycoside resistance in mycobacterium tuberculosis. Mol Microbiol 77:830–840.

11. Gagneux S, Long CD, Small PM, Van T, Schoolnik GK, Bohannan BJM. 2006. The Competitive Cost of Antibiotic Resistance in Mycobacterium tuberculosis. Science (80-) 312:1944–1946.

12. Sander P, Springer B, Prammananan T, Sturmfels A, Kappler M, Pletschette M, Böttger EC. 2002. Fitness cost of chromosomal drug resistance-conferring mutations. Antimicrob Agents Chemother 46:1204–1211.

13. Dye C, Williams BG, Espinal MA, Raviglione MC. 2002. Erasing the world’s slow stain: strategies to beat multidrug-resistant tuberculosis. Science (80-) 295:2042–2046.

14. Cohen T, Dye C, Colijn C, Williams B, Murray M. 2009. Mathematical models of the epidemiology and control of drug-resistant TB. Expert Rev Respir Med 3:67–79.

15. Borrell S, Gagneux S. 2009. Infectiousness, reproductive fitness and evolution of drug-resistant Mycobacterium tuberculosis. Int J Tuberc Lung Dis 13:1456–1466.

16. Mesfin YM, Hailemariam D, Biadglign S, Kibret KT. 2014. Association between HIV/AIDS and multi-drug resistance tuberculosis: A systematic review and meta-analysis. PLoS One 9:1–9.

17. Suchindran S, Brouwer ES, Van Rie A. 2009. Is HIV infection a risk factor for multi-drug resistant tuberculosis? A systematic review. PLoS One 4.

18. Khan PY, Yates TA, Osman M, Warren RM, van der Heijden Y, Padayatchi N, Nardell EA, Moore D, Mathema B, Gandhi N, Eldholm V, Dheda K, Hesseling AC, Mizrahi V, Rustomjee R, Pym A. 2019. Transmission of drug-resistant tuberculosis in HIV-endemic settings. Lancet Infect Dis 19:e77–e88.

19. Ssengooba W, Lukoye D, Meehan CJ, Kateete DP, Joloba ML, De Jong BC, Cobelens FG, Van Leth F. 2017. Tuberculosis resistance-conferring mutations with fitness cost among HIV-positive individuals in Uganda. Int J Tuberc Lung Dis 21:531–536.

20. Sander P, Springer B, Prammananan T, Sturmfels A, Kappler M, Pletschette M, Böttger EC. 2002. Fitness cost of chromosomal drug resistance-conferring mutations. Antimicrob Agents Chemother 46:1204–1211.

21. Billington OJ, Mchugh TD, Gillespie SH. 1999. Physiological cost of rifampin resistance induced in vitro in Mycobacterium tuberculosis. Antimicrob Agents Chemother 43:1866–1869.

22. Egger M, Ekouevi DK, Williams C, Lyamuya RE, Mukumbi H, Braitstein P, Hartwell T, Graber C, Chi BH, Boulle A, Dabis F, Wools-Kaloustian K. 2012. Cohort profile: The international epidemiological databases to evaluate AIDS (IeDEA) in sub-Saharan Africa. Int J Epidemiol 41:1256–1264.

23. Mcgowan CC, Cahn P, Gotuzzo E, Padgett D, Pape JW, Wolff M, Schechter M, Masys DR. 2007. Cohort Profile: Caribbean, Central and South America Network for HIV research (CCASAnet) collaboration within the International Epidemiologic Databases to Evaluate AIDS (IeDEA) programme. Int J Epidemiol 36:969–976.

24. Gagneux S, Burgos M V., DeRiemer K, Enciso A, Muñoz S, Hopewell PC, Small PM, Pym AS. 2006. Impact of bacterial genetics on the transmission of isoniazid-resistant Mycobacterium tuberculosis. PLoS Pathog 2:0603–0610.

25. Pym AS, Saint-Joanis B, Cole ST. 2002. Effect of katG mutations on the virulence of Mycobacterium tuberculosis and the implication for transmission in humans. Infect Immun 70:4955–4960.

26. Heym B, Alzari PM, Honore N, Cole ST. 1995. Missense mutations in the catalsase-peroxidase gene, katG, are associated with isoniazid resistance in Mycobacterium tuberculosis. Mol Microbiol 15:235–245.

27. van Soolingen D, de Haas PEW, van Doorn HR, Kuijper E, Rinder H, Borgdorff MW. 2000. Mutations at Amino Acid Position 315 of the katG Gene Are Associated with High-Level Resistance to Isoniazid, Other Drug Resistance, and Successful Transmission of Mycobacterium tuberculosis in The Netherlands. J Infect Dis 182:1788–1790.

28. Billington OJ, McHugh TD, Gillespie SH. 1999. Physiological cost of rifampin resistance induced in vitro in Mycobacterium tuberculosis. Antimicrob Agents Chemother 43:1866–9.

29. Mariam DH, Mengistu Y, Hoffner SE, Andersson DI. 2004. Effect of rpoB Mutations Conferring Rifampin Resistance on Fitness of Mycobacterium tuberculosis. Antimicrob Agents Chemother 48:1289–1294.

30. Sandgren A, Strong M, Muthukrishnan P, Weiner BK, Church GM, Murray MB. 2009. Tuberculosis drug resistance mutation database. PLoS Med 6:0132–0136.

31. Zürcher K, Ballif M, Fenner L, Borrell S, Keller PM, Gnokoro J, Marcy O, Yotebieng M, Diero L, Carter EJ, Rockwood N, Wilkinson RJ, Cox H, Ezati N, Abimiku AG, Collantes J, Avihingsanon A, Kawkitinarong K, Reinhard M, Hömke R, Huebner R, Gagneux S, Böttger EC, Egger M. 2019. Drug susceptibility testing and mortality in patients treated for tuberculosis in high-burden countries: a multicentre cohort study. Lancet Infect Dis 19:298–307.

32. Kwan C, Ernst JD. 2011. HIV and tuberculosis: A deadly human syndemic. Clin Microbiol Rev 24:351–376.

33. Hanrahan CF, Theron G, Bassett J, Dheda K, Scott L, Stevens W, Sanne I, Van Rie A. 2014. Xpert MTB/RIF as a measure of sputum bacillary burden: Variation by HIV status and immunosuppression. Am J Respir Crit Care Med 189:1426–1434.

34. Casali N, Nikolayevskyy V, Balabanova Y, Harris SR, Ignatyeva O, Kontsevaya I, Corander J, Bryant J, Parkhill J, Nejentsev S, Horstmann RD, Brown T, Drobniewski F. 2014. Evolution and transmission of drug-resistant tuberculosis in a Russian population. Nat Genet 46:279–286.

35. Eldholm V, Monteserin J, Rieux A, Lopez B, Sobkowiak B, Ritacco V, Balloux F. 2015. Four decades of transmission of a multidrug-resistant Mycobacterium tuberculosis outbreak strain. Nat Commun 6:7119.

36. Song T, Park Y, Shamputa IC, Seo S, Lee SY, Jeon HS, Choi H, Lee M, Glynne RJ, Barnes SW, Walker JR, Batalov S, Yusim K, Feng S, Tung CS, Theiler J, Via LE, Boshoff HIM, Murakami KS, Korber B, Barry CE, Cho SN. 2014. Fitness costs of rifampicin resistance in Mycobacterium tuberculosis are amplified under conditions of nutrient starvation and compensated by mutation in the β′ subunit of RNA polymerase. Mol Microbiol 91:1106–1119.

37. Vogwill T, Kojadinovic M, MacLean RC. 2016. Epistasis between antibiotic resistance mutations and genetic background shape the fitness effect of resistance across species of Pseudomonas. Proceedings Biol Sci 283.

38. Castro RAD, Ross A, Kamwela L, Reinhard M, Loiseau C, Feldmann J, Borrell S, Trauner A, Gagneux S. 2019. The Genetic Background Modulates the Evolution of Fluoroquinolone-Resistance in Mycobacterium tuberculosis. Mol Biol Evol 37:195–207.

39. Huang CC, Tchetgen ET, Becerra MC, Cohen T, Hughes KC, Zhang Z, Calderon R, Yataco R, Contreras C, Galea J, Lecca L, Murray M. 2014. The effect of HIV-related immunosuppression on the risk of tuberculosis transmission to household contacts. Clin Infect Dis 58:765–774.

40. Wells CD, Cegielski JP, Nelson LJ, Laserson KF, Holtz TH, Finlay A, Castro KG, Weyer K. 2007. HIV Infection and Multidrug-Resistant Tuberculosis—The Perfect Storm. J Infect Dis 196:S86–S107.

